# Global phylogeography of *Amrasca biguttula* (Hemiptera: Cicadellidae) across eight countries reveals a single-haplotype incursion into the United States beyond its putative native range

**DOI:** 10.64898/2025.12.31.697166

**Authors:** Muhammad Z. Ahmed, Nisha Yadav, Sachin Rustgi, Gautam Saripalli, Isaac L. Esquivel, Tim B. Bryant, Scott Graham, Alana L. Jacobson, Midhula Gireesh, Shimat V. Joseph, Alejandro Del Pozo-Valdivia, Karla M. Addesso, Tyler J. Raszick, Rafia Khan, Jose C. V. Rodrigues, Tom R. Bilbo, Jeremy K. Greene, Francis F. P. Reay-Jones

## Abstract

*Amrasca biguttula* is a pest of cotton, ornamentals, and vegetables, and is invading the southeastern United States. We addressed five questions: (1) Are US populations genetically identical to those in the putative native range? (2) Does the southeastern US outbreak extend the recent Puerto Rico and Florida invasion or represent separate introductions? (3) Does the lineage infesting ornamentals match that on other crops? (4) What is the mitochondrial cytochrome oxidase subunit 1 (*mtCOI*) diversity of *A. biguttula*? (5) Which regions constitute its native versus invaded range? Adults were collected from Florida, Georgia, South Carolina, Texas, and Puerto Rico. Species identity was confirmed morphologically and by sequencing *mtCOI*, where possible, from five individuals per site, yielding consensus haplotypes. We analyzed 11 US, 20 Puerto Rican, and 342 sequences originating from publicly available databases (n = 373). Phylogenetic analysis confirmed monophyly of this species. A haplotype network recovered 70 unique haplotypes, including a star-like core of 68 radiating from Hap01 and two divergent singletons. The mainland US and Chinese populations were fixed for Hap01, indicating recent introductions. Puerto Rico exhibited Hap01, Hap52, and Hap70, reflecting multiple introductions, but could also result from haplotype diversity carried within a single introduction involving many individuals. Furthermore, southeastern US sites were fixed for Hap01, indicating extension from Puerto Rico and Florida. No haplotype was host-specific, confirming a lineage across crops. Diversity metrics indicate recent expansion in Iraq, Hong Kong, and Puerto Rico. South Asia, particularly Bangladesh, India, and Pakistan, showed the highest haplotype diversity, supporting its role as the putative native range. In contrast, low diversity in China and the US is consistent with recent invasion. Curation yielded 373 high-quality sequences, providing a unified database to support quarantine and management strategies.

## 1 INTRODUCTION

*Amrasca biguttula* (Ishida) (Hemiptera: Cicadellidae), commonly known as the two-spot leafhopper or cotton jassid, is a minute that feeds on the contents of parenchyma cells and causes significant yield losses on infested plants across South and Southeast Asia (Sagarbarria et al., 2020). *Amrasca biguttula*, historically known as the “Indian cotton jassid” on cotton (*Gossypium* spp.) (Merino, 1936), has since expanded its host range and geographic distribution, infesting cotton, ornamental plants, and vegetable crops in China, India, Iraq, and Pakistan (Kranthi et al., 2018; Jaod & Nawar, 2023; Sharma et al., 2025). Feeding by nymphs and adults produces the characteristic “hopperburn,” marked by necrotic leaf margins and reduced photosynthetic area lead to substantial yield reductions. In eggplant (*Solanum melongena* L.), losses may reach 37% or more as leafhopper populations increase (Ahmed, 1982); in okra (*Abelmoschus esculentus* (L.) Moench), up to 50% (Devi et al., 2018); and in cotton (*Gossypium hirsutum* L.), a 37% loss has been recorded (Ahmad et al., 1985), with overall reductions of 25–45% (Bhat et al., 1986). Severe infestations may lead to outbreaks of secondary pests (Rozas et al., 2017) and can also facilitate the transmission of phytoplasmas and viruses (Al-Hamdany & Al-Karboli, 2017). *Amrasca biguttula* is a highly polyphagous leafhopper, inflicting damage on okra, eggplant, peanut (*Arachis hypogaea* L.), sunflower (*Helianthus annuus* L.), roselle (*Hibiscus sabdariffa* L.), and numerous ornamentals (Chupungco et al., 2014; Saeed et al., 2015). Among these, ornamental hibiscus poses a unique risk: our recent survey in South Carolina detected infestations on nursery-grown plants adjacent to cotton and vegetable fields (Ahmed MZ, pers. obs.), and hopperburn damage has been confirmed in Puerto Rico (Liburd et al., 2024). Because hibiscus is promoted as a pollinator-friendly ornamental, growers often avoid insecticide applications on blooms, and “pollinator-safe, no-spray” labeling further increases its market appeal (Khachatryan & Rihn, 2016). This combination poses a dual challenge: untreated, infested hibiscus may be moved across or out of state lines, triggering regulatory scrutiny and potential quarantine measures (Liburd et al., 2024); meanwhile, year-round foliage in landscapes provides a refuge for overwintering, leading to early-season population buildups of *A. biguttula* that threaten nearby crops, including cotton and vegetables.

The control of *A. biguttula* is complicated by the rapid development of resistance to neonicotinoid, organophosphate, and pyrethroid insecticides, particularly in Pakistan and India (Abbas et al., 2018; Akmal et al., 2020; Sharma et al., 2025). Although integrated pest management strategies, including endophytic and foliar applications of *Beauveria bassiana* and *Metarhizium* spp., offer promising alternatives, their field efficacy against *A. biguttula* remains underexplored (Allegrucci et al., 2017; Rozas et al., 2018).

Recent records demonstrate that *A. biguttula* is expanding beyond its putative native range. In China, it infests cotton and okra (Sharma et al., 2025), while in Iraq, it has emerged on cowpea (*Vigna unguiculata* (L.) Walp.), eggplant, and okra (Al-Hamdany & Al-Karboli, 2017; Jaod & Nawar, 2023). In 2023, *A. biguttula* was detected for the first time in the Western Hemisphere, attacking both cultivated and wild cotton, eggplant, and hibiscus in Puerto Rico (Cabrera-Asencio et al., 2023). A single specimen from Cuba in 1956 suggested an earlier, undocumented introduction (Cabrera-Asencio et al., 2023), and its recent confirmation in Puerto Rico (Cabrera-Asencio et al., 2023) and Florida (Liburd et al., 2024) has heightened concerns about the potential further spread throughout the southeastern United States, including Alabama, Georgia, Louisiana, Mississippi, North Carolina, South Carolina and Texas (Ahmed, 2025a; Esquivel et al., 2025). In addition, in 2024, it was reported on cotton in Central Africa (Cameroon) by Jacques et al. (2024) and on okra and roselle in West Africa (Niger) by Akonde et al. (2024). These observations raise several critical questions about the invasion dynamics of *A. biguttula*: are US populations genetically identical to those in the putative native range; do South Carolina and southeastern US outbreaks reflect northward spread from Florida or independent introductions; are different lineages associated with different host plants; and how does global *mtCOI* diversity delineate native versus invaded regions?

Morphological identification of *A. biguttula* is challenging because immature instars are difficult to distinguish, and adult diagnosis requires detailed examination of male genitalia and pregenital tergites (Xu et al., 2017). Molecular analysis of the *mtCOI* barcode region has therefore become an indispensable complement to morphology, enabling the reliable confirmation of species, discrimination of haplotypes, and reconstruction of invasion pathways (Hebert et al., 2003; Kranthi et al., 2018). Although nuclear markers such as 18S and 28S rRNA genes (Buckman et al., 2013) and internal transcribed spacers (Rugman-Jones et al., 2006) have been developed for the identification of insect pests, *mtCOI* remains the definitive molecular marker for genetic identification and population genetic analyses due to its high nucleotide substitution rate.

To date, *mtCOI* studies of *A. biguttula* have primarily focused on regional populations in Pakistan (Akmal et al., 2017), Bangladesh (Hossain et al., 2023), Iraq (Jaod & Nawar, 2023), India (Kranthi et al., 2017; Sharma et al., 2025), and the Philippines (Sagarbarria et al., 2020). However, these studiesare limited by host range and narrow geographic scope, underscoring the need for broader, more integrative analyses. Variations in primer design, fragment length, and annotation standards have precluded direct cross-study comparisons. Here, we curate 373 *mtCOI* sequences from diverse sources, uniformly trim them to a standardized fragment, and collapse them into 70 consensus haplotypes. This global framework minimizes bias and enables robust assessment of worldwide genetic diversity, host-associated lineages, and invasion pathways for this invasive pest.

Our unified haplotype database will enable the rapid and accurate identification of global *A. biguttula* haplotypes, inform regulatory decisions, and underpin integrated pest management and future phylogeographic studies.

## 2 MATERIALS AND METHODS

### 2.1 Sample collection and DNA extraction

Adult *A. biguttula* were sampled from five host-associated populations across Florida (okra, *A. esculentus*), Georgia (hibiscus, *H. rosa-sinensis*; cotton, *G. hirsutum*), Puerto Rico (cotton, *G. hirsutum*; eggplant, *S. melongena*), South Carolina (hibiscus, *H. rosa-sinensis*; cotton, *G. hirsutum*; eggplant, *S. melongena*), and Texas (hibiscus, *H. rosa-sinensis*). At each site, up to ten adults were collected, and these individuals together constituted a single population (n = 10 per site; Table S1). Specimens were morphologically identified as *A. biguttula* based on tentative identification characters, including two distinct black forewing spots, body coloration, and size (Cabrera-Asencio et al., 2023). Genomic DNA was extracted individually from five randomly selected individuals per population, where possible (*n* = 25 in total across five populations), using the DNeasy Blood & Tissue Kit (Qiagen, MD, US), following the method described by Ahmed et al. (2024). For Puerto Rico samples, the DNA was isolated using the Promega Maxwell RSC Genomic DNA kit (Promega, WI, USA). Polymerase Chain Reaction (PCR) amplification and sequencing of the *mtCOI* sequence from each extract revealed identical haplotypes within populations, allowing us to retain one high-quality consensus sequence per population (NY05, NY13, NY22, NY30, and NY33; Table S1) for comparative analyses.

### 2.2 PCR amplification and sequencing

A ∼700 bp fragment of the *mtCOI* gene was amplified using universal primers LCO1490 (5′-GGT CAA CAA ATC ATA AAG ATA TTG G-3′) and HCO2198 (5′-TAA ACT TCA GGG TGA CCA AAA AAT CA-3′) (Folmer et al., 1994). Each 25 µL reaction comprised 12.5 µL of 2× Terra PCR Direct Buffer (Takarabio, CA, US) containing MgCl₂ and dNTPs, 0.8 µL of each primer (10 µM), 0.5 µL of Terra PCR Direct Polymerase Mix (Takarabio, CA, US), 4.0 µL of genomic DNA (10–50 ng), and nuclease-free water to final volume. Thermal cycling was performed on a Bio-Rad C1000 (Bio-Rad, CA, US) Touch Thermal Cycler with the following program: initial denaturation at 98 °C for 2 min; 40 cycles of 98 °C for 10 s, 54 °C for 15 s, and 68 °C for 1 min; a final extension at 68 °C for 5 min; and a hold at 4 °C.

PCR products were resolved on 1.5% agarose gels stained with ethidium bromide and visualized under ultraviolet illumination. Amplicons of the expected size were purified and submitted to Molecular Cloning Laboratories (MCLAB, CA, US) for bidirectional Sanger sequencing using the same primer pair. Samples collected in Puerto Rico were amplified and sequenced at the USDA-APHIS-PPQ Science & Technology, Insect Management and Molecular Diagnostics Laboratory, Edinburg, Texas, USA. Sequencing reads were assembled, trimmed, and manually inspected in BioEdit v7.2.5 (Hall, 1999).

### 2.3 Sequence curation, length-based grouping, alignment, and translation

In addition to our 31 newly generated sequences, we retrieved 364 *A. biguttula mtCOI* records from GenBank and the Barcode of Life Database (BOLD), resulting in an initial dataset of 395 sequences (Table S1). These records ranged from 356 bp (KU684386; Akmal et al., 2017) to 701 bp (MK293719; Ahmad, 2019), with the modal length of 487 bp represented by 97 accessions (KX813719–KX813815; Kranthi et al., 2017). Twenty-two sequences were excluded due to insufficient length (see Table S1). Because these sequences originated from multiple laboratories using different primer sets, PCR conditions, and sequencing platforms, standardizing on a common 487 bp *mtCOI* fragment provided consistent taxon coverage while preserving the 95 polymorphic sites essential for haplotype-network reconstruction and diversity inference. For descriptive purposes, the untrimmed sequences fell into four length categories: 487 bp, 500–599 bp, 600–699 bp, and 700 bp.

All 373 sequences —representing four untrimmed length categories (487 bp, 500–599 bp, 600–699 bp, and 700 bp)—were aligned using ClustalW v2.1 (Thompson et al., 1994) implemented in MEGA11 v11.0.13 (Tamura et al., 2021). Alignments were translated using MEGA11 v11.0.13 to confirm the presence of intact open reading frames and the absence of stop codons. Although longer alignments (Groups 2–4) contributed additional singleton polymorphisms, trimming all sequences to a 487 bp fragment (Group 1) preserved the vast majority of informative sites while eliminating gapped positions relative to shorter reads and provided 100% data coverage.

### 2.4 Haplotype assignment and genetic analysis

We performed all quality-control and translation checks in R (R Core Team, 2024) to ensure our 487 bp *mtCOI* alignment was free of indels, ambiguous bases, and stop codons. The FASTA alignment was read into R using the seqinr package (Charif & Lobry, 2007), which allowed us to scan each sequence for gaps (“–”) and ambiguous “N” characters and to verify that every sequence length was a multiple of three. We then translated all sequences under the invertebrate mitochondrial genetic code (NCBI code 5) across all six reading-frame/strand combinations. This screening identified the forward strand with a two-nucleotide offset (frame 2) as yielding zero internal stop codons. Final stop-codon confirmation and amino-acid sequence extraction were performed using the Biostrings package (Pagès et al., 2024).

Haplotype assignment was performed using four complementary approaches: Kimura 2-parameter genetic distances (Kimura, 1980), statistical parsimony in TCS v1.23 (Clement et al., 2000), median-joining networks in PopART v1.7 (Leigh & Bryant, 2015), and polymorphism analysis in DnaSP v6.12.03 (Rozas et al., 2017). Genetic distances and substitution patterns were estimated using the Maximum Composite Likelihood model (Tamura et al., 2004), incorporating all codon positions (1^st^, 2^nd^ and 3^rd^) as well as noncoding regions. Ambiguous positions were removed by pairwise deletion, and the final dataset included 487 positions across 70 unique haplotypes (Table S1). The nucleotide composition and Kimura 2-parameter (K2P) genetic distance (Kimura, 1980) between haplotypes were calculated in MEGA11 v11.0.13 (Tamura et al., 2021). We calculated nucleotide diversity using DnaSP v5.0 (Librado & Rozas, 2009) and poprat-1.7 (Leigh & Bryant, 2015). The pairwise Φst (*Fst*) values were calculated to identify limits to gene flow among these haplotypes (Excoffier et al., 1992). Analysis of molecular variance (AMOVA) was undertaken with 1,000 permutations using poprat-1.7 (Leigh & Bryant, 2015). We also conducted neutrality tests (Fu’s Fs and Tajima’s D) using DnaSP v5.0 to explore the possibility of the recent population expansion in *A. biguttula* (Librado & Rozas, 2009). Tajima’s D was calculated on the aligned sequence dataset using DnaSP v5.0 with 10,000 coalescent simulations to assess the deviation from neutral expectations.

### 2.5 Phylogenetic analysis

Although haplotype analysis of 373 *mtCOI* sequences yielded 70 unique *A. biguttula* haplotypes, phylogenetic reconstruction incorporated all 373 sequences plus 24 outgroup taxa representing Empoascini, Dikraneurini, and Erythroneurini lineages (Figure 1). The sequences were aligned using MUSCLE with default parameters, resulting in a final dataset of 487 nucleotide positions using the aforementioned approach. In our phylogenetic analysis, the final dataset comprised 70 unique haplotypes alongside 24 outgroup sequences (94 sequences in total). Ambiguously aligned regions were manually inspected and trimmed to ensure positional homology across all taxa. Phylogenetic reconstruction was performed using IQ-TREE v1.6.12 (Nguyen et al., 2015) based on a multiple-sequence alignment of 94 sequences, including 70 unique haplotypes and 24 outgroup taxa spanning 487 nucleotide sites. ModelFinder (Kalyaanamoorthy et al., 2017) identified K3Pu+F+G4 as the best-fit substitution model under the Bayesian Information Criterion. This model employs unequal purine transition rates (K3Pu), empirical base frequencies (+F), and gamma-distributed rate heterogeneity across four categories (+G4), with an estimated shape parameter α = 0.3543. Node support was evaluated via ultrafast bootstrap approximation (UFBoot2) with 10,000 replicates (Hoang et al., 2017). The maximum-likelihood tree achieved a log-likelihood of –3632.437, a total branch length of 1.6117, and internal branches totaling 0.8325 (51.65 % of the tree length). A consensus tree was built from the 10,000 bootstrap replicates, optimizing branch lengths on the original alignment and retaining all nodes with nonzero support.

**FIGURE 1.**
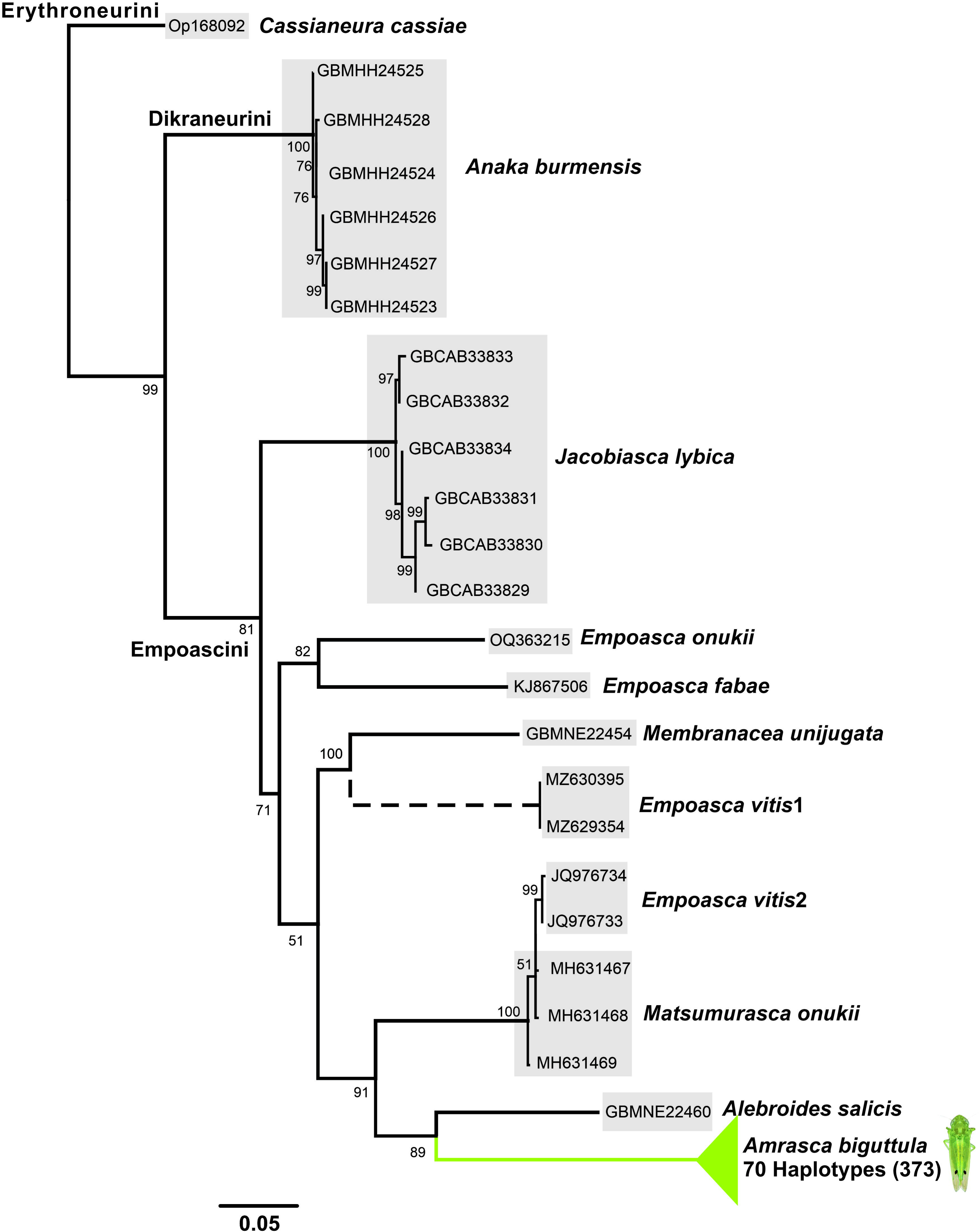
Maximum likelihood phylogeny of *Amrasca biguttula* and related taxa inferred using IQ-TREE v1.6.11 under the best-fit substitution model K3Pu+F+G4, selected via ModelFinder. Node support values represent ultrafast bootstrap (UFBoot2) percentages based on 10,000 replicates. Branch lengths are scaled to substitutions per site. The tree includes 70 haplotypes of *A. biguttula* and 24 outgroup sequences, with *Cassianeura cassiae* designated as the primary outgroup based on Alagarsamy et al. et al. (2024). Additional outgroup taxa include *Empoasca vitis* (4), *Empoasca onukii* (1), *Matsumurasca onukii* (3), *Alebroides salicis* (1), *Membranacea unijugata* (1), *Jacobiasca lybica* (6), *Anakaburmensis* (6), and *Empoasca fabae* (1). The hatched line denotes a subclade that falls outside the main *Empoasca vitis* 2 clade; its placement is unresolved with *mtCOI* alone and may require additional genes or expanded sampling to resolve.

Guided by the phylogenetic framework of Alagarsamy et al. (2024), we designated *Cassianeura cassia* as the primary outgroup (1 sequence) based on its basal position in Erythroneurini (see their Fig. 3). To encompass the major Empoascini, Dikraneurini, and Erythroneurini lineages, we then included *Empoasca vitis* (4 sequences), *Empoasca onukii* (1), *Matsumurasca onukii* (3), *Alebroides salicis* (1), *Membranacea unijugata* (1), *Jacobiasca lybica* (6), *Anaka burmensis* (6), and *Empoasca fabae* (1). These taxa collectively span the phylogenetic breadth of the three tribes, providing a robust scaffold for rooting the tree and accurately interpreting the placements of all 373 *A. biguttula* sequences. At least three recent phylogenomic and mitogenomic studies recover Empoascini and Dikraneurini as a strongly supported sister-group clade to Erythroneurini using maximum-likelihood and Bayesian inference, thus justifying our choice of Erythroneurini as the outgroup in this study (Cao et al., 2023; Chen et al., 2021; Lin et al., 2021).

### 2.6 Network construction and outgroup inclusion

Genealogical relationships among haplotypes were inferred using statistical parsimony and median-joining methods. Parsimony networks were constructed using the algorithm of Templeton et al. (1992), with a 95% connection threshold, and gaps were treated as a fifth character state (De Barro & Ahmed, 2011). Median-joining networks were generated in PopART v1.7 with epsilon = 0 and 5,000 iterations.

### 2.7 Data analysis

Nucleotide diversity per site (π) for each population was estimated by non-parametric bootstrap resampling of aligned *mtCOI* sequences. For each population, 1,000 bootstrap replicates were generated by sampling alignment columns with replacement, and nucleotide diversity (π) was recalculated for each replicate. These bootstrap distributions of π served as independent observations for downstream testing. We compared median π across eight populations (Overall, Bangladesh, Hong Kong, India, Iraq, Pakistan, Philippines, and Puerto Rico) using the Kruskal–Wallis rank-sum test (α = 0.05). Following a significant Kruskal–Wallis result, we performed pairwise Dunn’s tests with Benjamini–Hochberg correction to identify which population pairs differed. Populations were assigned group letters so that any two sharing a letter do not differ at p < 0.05. All analyses were conducted in R v 4.1.2 with the boot, FSA, and pgirmess packages. We quantified, for each of the 70 haplotypes, the number of sequences (frequency), associated host-plant species, host-plant families, and unique countries of occurrence (Table S2). Summary tables were generated in R v4.1.2 to characterize host-plant association and geographic breadth across haplotypes.

To test whether haplotype prevalence was associated with host-plant association or geographic breadth, we performed Spearman’s rank correlation analyses between sequence count and three variables: host-plant species richness, host-plant family richness, and country count. All tests were two-tailed, conducted using the *cor.test()* function in R v4.1.2, with a significance threshold of α = 0.05.

## 3 RESULTS

### 3.1 Single *mtCOI* haplotype across US *A. biguttula* populations

All specimens were morphologically confirmed as *A. biguttula* based on the presence of two distinct black spots on the forewings, overall body coloration, and diagnostic size. Genomic DNA, where possible, was extracted from five individuals per population (total n = 25), yielding high-quality *mtCOI* amplifications. Within each population, all five sequences were identical; therefore, a single 577 bp consensus sequence was retained for each site (NY05, NY13, NY22, NY30, and NY33) for downstream analyses. BLASTn searches against publicly available *mtCOI* entries in GenBank and The Barcode of Life Data Systems (BOLD) revealed that all US populations by host plants matched with 100% identity to GenBank accession KU684356 (designated as Hap01 in this study), with zero nucleotide differences over the whole 577 bp fragment. Pairwise distance analysis further confirmed zero p-distance among the consensus sequences of all US population representatives and KU684356, indicating a single haplotype across all samples. No nucleotide polymorphism was detected among host plants (cotton, eggplant, hibiscus, okra) or among continental geographic regions (Florida, Georgia, South Carolina, and Texas). However, three distinct haplotypes were identified in Puerto Rico, indicating localized genetic variation. These results demonstrate that all newly sequenced *A. biguttula* specimens share a single *mtCOI* haplotype, identical to KU684356, with no evidence of intraspecific variation or possible cryptic speciation (as observed in other invasive species) across hosts or states in these samples.

### 3.2 Consensus sequences in a 373-sequence *mtCOI* global dataset

To assess broader intraspecific variation, each consensus sequence was analyzed together with 342 publicly available *mtCOI* entries in GenBank and BOLD, yielding a total of 373 aligned sequences. Using the Kimura 2-parameter model for genetic distances, statistical parsimony, median-joining network construction, and polymorphism analysis, we recovered 70 unique *mtCOI* haplotypes. These included one dominant haplotype, Hap01 (n = 239; 64.1%), a second Hap02 (n = 8; 2.1%), 19 mid-frequency haplotypes represented by 73 individuals (19.6%), 9 low-frequency haplotypes represented by 12 individuals (3.2%), and 41 singletons (11.0%). This curated set of 70 haplotypes is presented here for the first time as a standardized reference framework for future phylogeographic and diagnostic studies (see Table S1).

### 3.3 Haplotype divergence

All *mtCOI* sequences aligned unambiguously over the trimmed 487 bp region, with no gaps, indels, or ambiguous characters. Each sequence was divided into three and translated under the invertebrate mitochondrial code in frame two without internal stop codons, confirming intact open reading frames in both R v4.1.2 and MEGA11. The haplotype dataset (70 unique haplotypes representing 373 sequences) showed markedly higher diversity. Across the 487 bp alignment, 95 sites were polymorphic, corresponding to 105 mutations (Eta). Each sequence corresponded to a unique haplotype (h = 70), yielding maximal haplotype diversity (Hd = 1.000; variance = 0.00001; SD = 0.002), indicating a highly diverse population. Nucleotide diversity was elevated (π = 0.01140 ± 0.00113; πJC = 0.01153), with an average of k = 5.552 nucleotide differences. Estimates of θ per site were 0.04474 (from Eta) and 0.04048 (Watterson’s θ), while θ per sequence reached 19.72. Under a finite-sites model, θ ranged from 0.01158 (from π) to 0.04728 (from Eta), consistently indicating high genetic variability. Phylogenetic reconstruction in IQ-TREE v1.6.12, based on the 487 bp alignment of 70 haplotypes plus 24 outgroups, identified K3Pu+F+G4 as the best-fit substitution model under BIC, with empirical base frequencies (A = 0.25, C = 0.152, G = 0.150, T = 0.448) and strong transition bias (A↔G, C↔T). The maximum likelihood tree (logL = –3632.44) had a total branch length of 1.6117, with ∼52% attributable to internal branches; however, 68 internal branches were near zero in length, reflecting shallow divergence among haplotypes. Collectively, these results confirm that the dataset comprises intact protein-coding *mtCOI* fragments, with the larger sequence set showing modest overall diversity and the haplotype-only dataset revealing maximal haplotype richness and elevated nucleotide diversity. Together with the shallow but structured phylogeny, these findings highlight both extensive standing variation and rapid diversification dynamics within the sampled populations.

### 3.4 Haplotype geographic substructure analysis

Geographic partitioning revealed distinct diversity profiles across countries, which were subsequently grouped into categories based on mitochondrial DNA diversity, reflecting their underlying demographic histories. Category 1 encompassed populations with low haplotype diversity (Hd) and low nucleotide diversity (π), consistent with recent bottlenecks or founder events; Category 2 included those with high Hd but low π, suggesting rapid expansion following a bottleneck; and Category 3 represented populations with both high Hd and high π, indicative of long-term evolutionary persistence and demographic stability (Table 2). Across all sequences (*n =* 373), spanning Bangladesh, China, Hong Kong, India, Iraq, Pakistan, the Philippines, Puerto Rico, and the US, we recovered h = 70 haplotypes, π = 0.01140 ± 0.00113, k = 5.552, and Hd = 1.000 ± 0.002 (Group a). Bangladesh (*n =* 4) exhibited h = 4, π = 0.01027 ± 0.00268, k = 5.000, and Hd = 1.000 ± 0.177, consistent with a large, stable population (Category 3; Group a). China (*n =* 1) and the US (*n =* 5) each yielded a single haplotype (Category 1), reflecting founder events with no measurable diversity. Hong Kong (*n =* 6) showed h = 2, π = 0.00205 ± 0.00103, k = 1.000, and Hd = 1.000 ± 0.500, consistent with rapid expansion after a bottleneck (Category 2; Group c). India (*n =* 239) harbored 43 haplotypes, π = 0.01213 ± 0.00159, k = 5.908, and Hd = 1.000 ± 0.005, reflecting long-term demographic stability (Category 3; Group a). Iraq (*n =* 8) exhibited two haplotypes, with π = 0.00821 ± 0.00411, k = 4.000, and Hd = 1.000 ± 0.500, consistent with reduced but non-trivial variation (Category 2; Group bc). Pakistan (*n =* 73) had 19 haplotypes, π = 0.00909 ± 0.00151, k = 4.427, and Hd = 1.000 ± 0.017, placing it among the high-diversity populations (Category 3; Group ab). The Philippines (*n =* 11) supported 10 haplotypes, π = 0.00370 ± 0.00036, k = 1.800, and Hd = 1.000 ± 0.045, consistent with rapid expansion after a bottleneck (Category 2; Group c). Puerto Rico (*n =* 20) yielded three haplotypes, π = 0.00274 ± 0.00091, k = 1.333, and Hd = 1.000 ± 0.272, also reflecting a bottleneck-expansion dynamic (Category 2; Group c).

Nucleotide diversity differed significantly among the eight populations (Kruskal–Wallis χ²₇ = 42.56, *p* < 0.0001; Table 1). Dunn’s post-hoc tests with Benjamini–Hochberg correction resolved four statistically distinct clusters that capture the major patterns of mitochondrial variation (Figure S1, Table S4).

**Table 1.** Genetic diversity parameters of *Amrasca biguttula* according to geographical locations.

| Haplotypes | Countries | h | Hp<sup>1</sup>(Hp<sup>2</sup>) | $\pi \pm SD$ | k | Hd ± SD |
| --- | --- | --- | --- | --- | --- | --- |
| Overall | Bangladesh (4), China (1), Hong Kong (6), India (239), Iraq (8), Pakistan (73), Philippines (11), Puerto Rico (20), US (11) | 373 | 70 (70) | 0.01140<sup>a</sup>± 0.00113 | 5.552 | 1.000± 0.002 |
| Hap01, Hap62–64 | Bangladesh**** | 4 | 4 (4) | 0.01027<sup>a</sup>± 0.00268 | 5.000 | 1.000± 0.177 |
| Hap01 | China* | 1 | 1 (2) | NA | NA | NA |
| Hap01, Hap66 | Hong Kong** | 6 | 2 (2) | 0.00205<sup>c</sup>± 0.00103 | 1.000 | 1.000± 0.500 |
| Hap01–13, Hap15–31, Hap33, 36, 41, 47–49, 53, 57–59, Hap67–69 | India**** | 239 | 43 (43) | 0.01213<sup>a</sup>± 0.00159 | 5.908 | 1.000± 0.005 |
| Hap01, Hap65 | Iraq** | 8 | 2 (2) | 0.00821<sup>bc</sup>± 0.00411 | 4.000 | 1.000± 0.500 |
| Hap01, 2, 6, 8, 14, 32, 34, 35, Hap37–40, Hap42–46, Hap60, 61 | Pakistan*** | 73 | 19 (19) | 0.00909<sup>ab</sup>± 0.00151 | 4.427 | 1.000± 0.017 |
| Hap01, 33, 37, 50–56 | Philippines** | 11 | 10 (10) | 0.00370<sup>c</sup>± 0.00036 | 1.800 | 1.000± 0.045 |
| Hap01, Hap52, Hap70 | Puerto Rico | 20 | 3(3) | 0.00274<sup>c</sup>± 0.00091 | 1.333 | 1.000± 0.272 |
| Hap01 | US* | 11 | 1 (1) | NA | NA | NA |
| h = total number of haplotype records |  |  |  |  |  |  |
| Hp<sup>1</sup> = count of distinct haplotypes inferred from parsimony-based and median-joining network analyses (using TCS v1.21 and PopART v1.7) |  |  |  |  |  |  |
| Hp<sup>2</sup> = number of unique haplotypes estimated from nucleotide diversity metrics (via DnaSP v5) |  |  |  |  |  |  |
| Hd = haplotype (gene) diversity |  |  |  |  |  |  |
| $\pi$ = nucleotide diversity per site | | | | | | |
| k = average number of nucleotide differences |  |  |  |  |  |  |
| Countries were grouped into diversity categories based on mitochondrial DNA variation: |  |  |  |  |  |  |
| *Category 1 — low Hd and low $\pi$, consistent with a recent bottleneck or founder event | | | | | | |
| **Category 2 — high Hd and low $\pi$, suggesting rapid expansion after a bottleneck | | | | | | |
| ***Category 3 — high Hd and high $\pi$, indicative of a large, stable population with long-term evolutionary persistence. These classifications reflect patterns of genetic structure commonly observed across geographically and demographically distinct populations. | | | | | | |
| Populations sharing the same letter are not significantly different at $p < 0.05$. | | | | | | |

### 3.5 Haplotype genetic distance

Overall, these 70 haplotypes in our dataset exhibited a mean pairwise genetic distance of 0.01169. When all self-comparisons (distance = 0) are excluded, inter-haplotype distances spanned from a minimum of 0.2055 (observed among multiple closely related pairs such as Hap03–Hap04, Hap14–Hap01, and Hap16–Hap01) to a maximum of 5.5515 (between Hap58 and Hap48), yielding a total range of 5.3460. The majority of haplotype pairs (approximately 65 %) differed by more than 1.0, underscoring substantial genetic structuring within our sample set and suggesting the coexistence of both shallow and deep population splits. Two haplotypes in particular, Hap48 (OP327267 from India) and Hap58 (KY012775 from India), fell outside the core network of interconnected sequences. Hap48 maintained a uniformly high minimal distance of 2.1146 to all other haplotypes and reached up to 3.8549 in its most divergent comparison. Hap58 was even more extreme, never less than 3.2266 divergent and peaking at 5.5515. Their pronounced genetic divergence, combined with clear isolation from the main haplotype cluster, suggests either long-term geographic separation or the existence of cryptic taxa. This pattern warrants targeted sampling and more comprehensive genomic analyses to resolve their evolutionary origins. Notably, no other haplotypes display similarly elevated divergence, reinforcing the likelihood that these two lineages reflect deeply separated populations or unrecognized taxonomic units. As a result, they disproportionately influence the overall genetic distance metrics within the dataset.

### 3.6 Phylogenetic analysis

To validate that the 70 *mtCOI* haplotypes, including 373 sequences from this study, published datasets (Akmal et al., 2017; Kranthi et al., 2017; Sagarbarria et al., 2020; Hossain et al., 2023; Jaod & Nawar, 2023; Sharma et al., 2025), and *mtCOI* entries retrieved from GenBank and BOLD belong to a single target species and lack cryptic diversity, we reconstructed a maximum-likelihood phylogeny rooted with appropriate outgroup taxa. The resulting tree recovered all haplotypes within a strongly supported monophyletic clade, confirming their conspecificity. Two haplotypes, Hap48 and Hap58, formed a distinct sister lineage to the main cluster, suggesting potential long-term isolation or unrecognized diversity that merits targeted sampling and deeper genomic investigation (Figure 1).

### 3.7 Genetic network and global diversity analysis

Statistical parsimony (95 % connection limit; maximum nine mutational steps; gaps as a fifth state) resolved three networks (Figure 2). A core network of 68 haplotypes radiates from Hap01 in a star-like topology. Hap01 has the highest outgroup weight (0.2656), with five secondary hubs: Hap07, Hap04, Hap05, Hap15 (each with a weight of 0.0415), and Hap03 (0.0332), which mark the most common mutational pathways. Singleton networks: Hap48 and Hap58 each form a separate, isolated network (weight = 1.0000 for both), reflecting extreme divergence. Parsimony probabilities remained high throughout the nine-step limit (*P*₉₅ % = 0.99921 at one step; 0.95471 at nine steps), confirming the network’s stability. The stark isolation of Hap48 and Hap58 suggests long-term separation or cryptic diversity, whereas the cohesive 68-haplotype core indicates recent diversification and substantial gene flow among the majority of lineages. We confirmed this structure with a median-joining network, which yielded an identical clustering pattern (Figure S2). The statistical parsimony network is shown in Figure 2. Haplotypes were grouped into haplogroups based on their frequencies and color-coded (cyan, light grey, light violet, magenta, orange, and red; Figs. 2 and 3) for AMOVA, which revealed significant differentiation among haplogroups (*Fst*, Φst = 0.05418, *p* < 0.001; Table 2).

**FIGURE 2.**
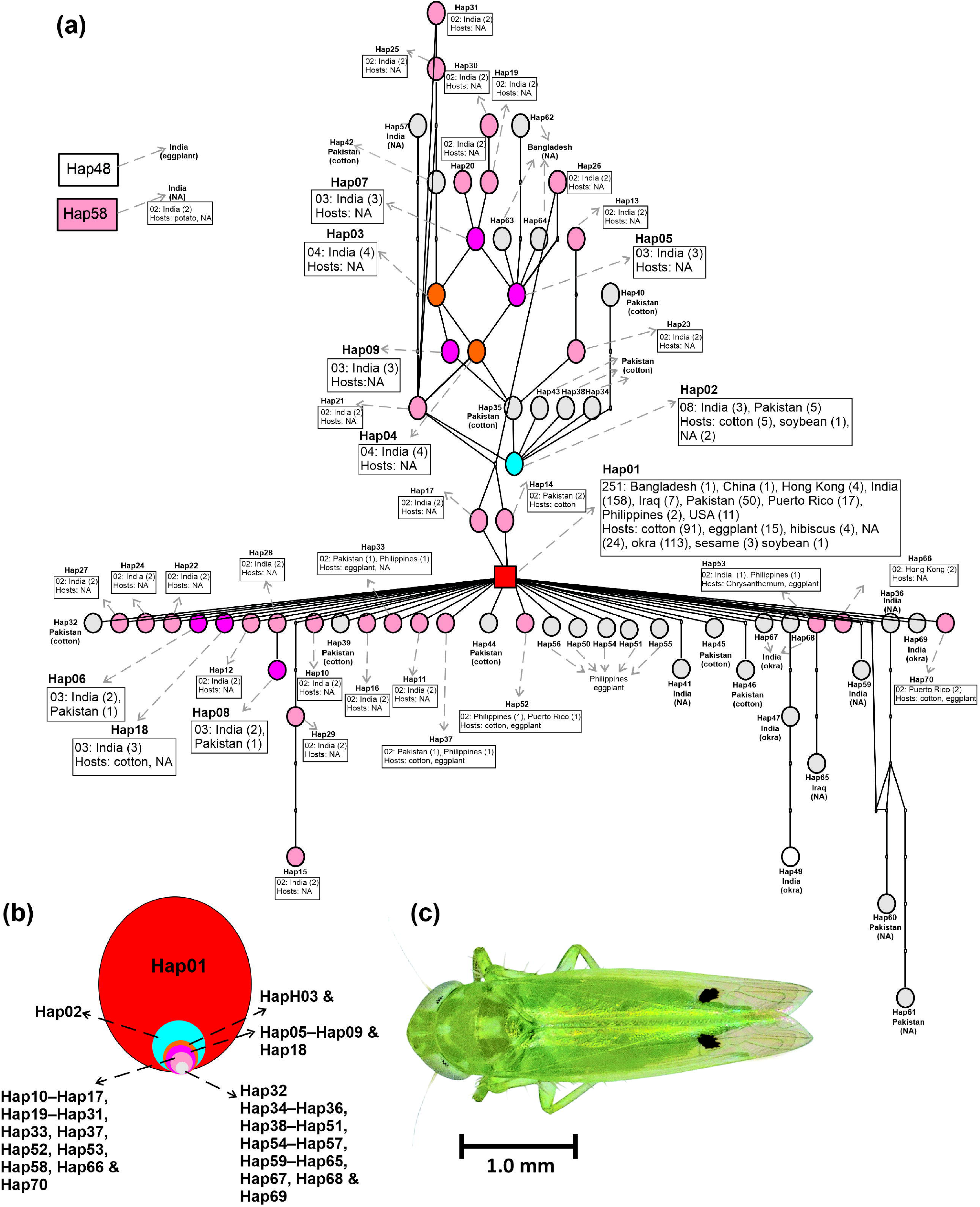
Genetic network analysis of 70 *mtCOI* sequences (487 bp) of *Amrasca biguttula* retrieved from GenBank and BOLD, based on consensus between statistical parsimony (a) (Templeton et al., 1992) implemented in TCS v.1.23 (Clement et al., 2000) and median-joining network method (Leigh & Bryant, 2015) in PopART v.1.7, at a 95% parsimony cutoff (maximum connection steps = 9; p(95%) = 0.9547) with gaps treated as a fifth state (De Barro & Ahmed, 2011). Please see Figure S2 for median-joining networks. Each oval denotes an individual haplotype (see Table S1). Small black circles represent inferred missing intermediates. Connections illustrate all plausible 95%-probability parsimonious links. The boxed number beside each node indicates the count of sequences sharing that haplotype (*n* > 1). Circle areas are proportional to the number of individuals in each haplotype or haplotype group (b). The adult of *A. biguttula* collected from a hibiscus host plant in Florence, South Carolina. A total of 70 haplotypes were identified and grouped by frequency into the following color-coded categories: red (Hap01, *n* = 1), cyan (Hap02, *n* = 1), orange (Hap03, Hap04, *n* = 2), magenta (Hap05–Hap09, Hap18, *n* = 6), light violet (Hap10–Hap17, Hap19–Hap31, Hap33, Hap37, Hap52, Hap53, Hap58, Hap66, Hap70, *n* = 28), and light grey (Hap32, Hap34–Hap36, Hap38–Hap51, Hap54–Hap57, Hap59–Hap65, Hap67–Hap69, *n* = 32).

**FIGURE 3.**
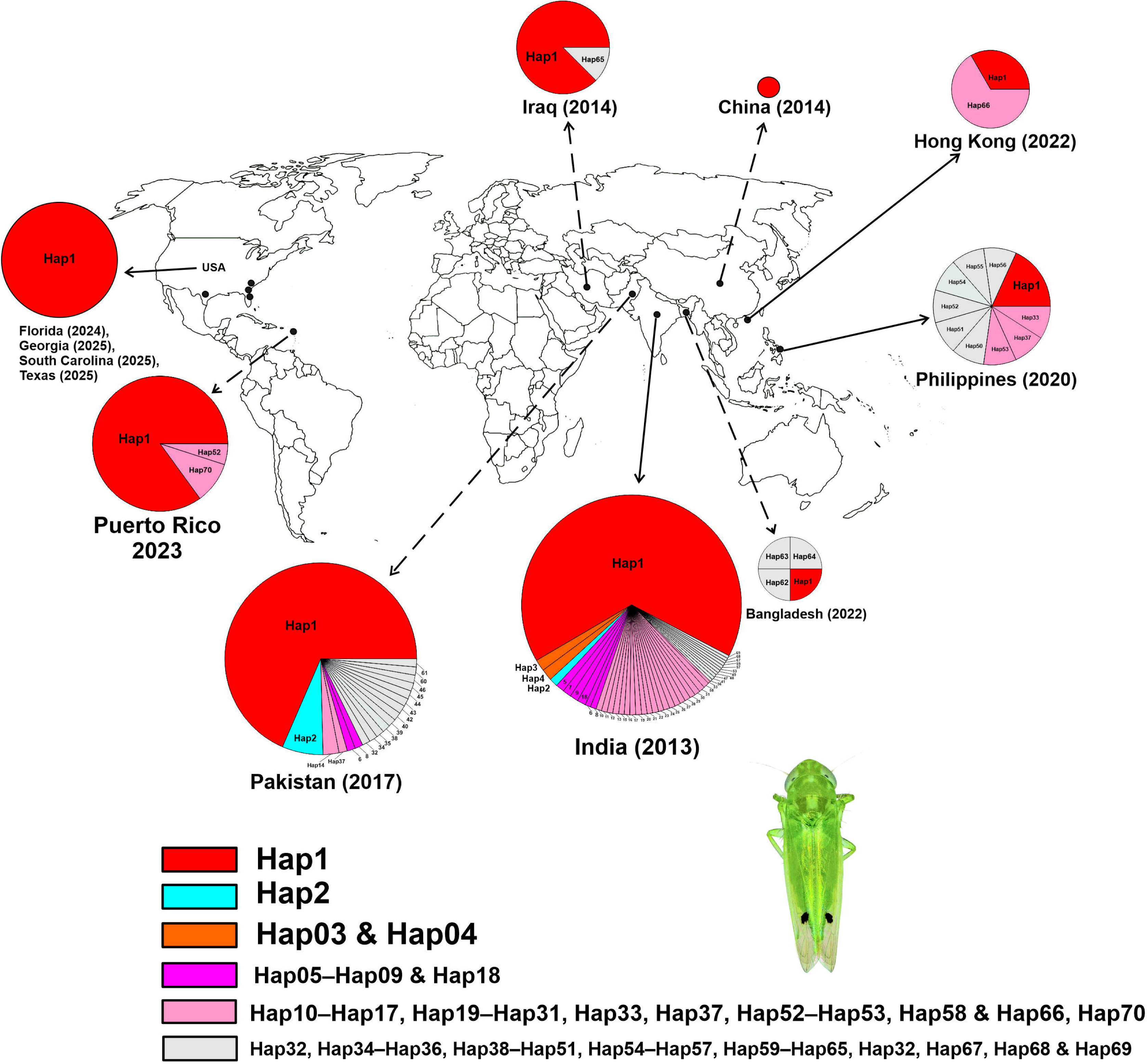
Global distribution of 70 *mtCOI* haplotypes of *Amrasca biguttula*. The year in parentheses indicates the first record of *A. biguttula* in each country, based on GenBank submissions or published reports. Circle areas are proportional to the number of sequences per haplotype, and colors denote abundance classes: the dominant haplotype Hap01 (*n* = 251 sequences); Hap02 (*n* = 8); Hap03 and Hap04 (*n* = 4 each); Hap05, Hap06, Hap07, Hap08, Hap09, and Hap18 (*n* = 3 each); Hap10–Hap17, Hap19–Hap31, Hap33, Hap37, Hap52, Hap53, Hap58, Hap66 and Hap70 (*n* = 2 each); and all remaining haplotypes (Hap32, Hap34–Hap36, Hap38–Hap45, Hap47–Hap51, Hap54–Hap57, Hap59–Hap65, Hap67–Hap69) represented by a single sequence (*n* = 1 each).

**Table 2.** Analysis of molecular variance (AMOVA) among haplotypes of *Amrasca biguttula* grouped by reported host plants and countries of origin.

| Source of Variation | df | Sum of Squares | Variance Components ($\sigma^2$) |
| --- | --- | --- | --- |
| **A. Haplotypes\*** |  |  |  |
| Among haplogroups# | 5 | 109.690 | 0.238 |
| Within haplogroups | 367 | 1526.846 | 4.160 |
| Total | 372 | 1636.536 | 4.399 |
| **B. Host Plants (Families)\*\*** |  |  |  |
| Among families | 5 | 129.046 | 0.291 |
| Within families | 367 | 1507.490 | 4.108 |
| Total | 372 | 1636.536 | 4.399 |
| **C. Countries\*\*\*** |  |  |  |
| Among countries## | 8 | 67.098 | 0.087 |
| Within countries | 364 | 1569.438 | 4.312 |
| Total | 372 | 1636.536 | 4.399 |
| **D. Continents\*\*\*\*** |  |  |  |
| Among continents | 1 | 1.604 | -0.158 |
| Within continents | 364 | 1569.438 | 4.312 |
| Total | 372 | 1636.536 | 4.358 |

\* $F_{st}$ , $\Phi_{st}$ = 0.05418, $p < 0.001$ , \*\* $F_{st}$ , $\Phi_{st}$ = 0.06614, $p = 0.004$ , \*\*\* $F_{st}$ , $\Phi_{st}$ = 0.01987, $p = 0.061$ \*\*\*\* $F_{st}$ , $\Phi_{st}$ = 0.01067, $p = 0.240$ (All values based on 1,000 permutations) Country-level variation was not statistically significant, likely due to #A total of 70 haplotypes were divided into groups based on their frequencies using colors (red, cyan, orange, magenta, light violet, and light grey) for AMOVA analysis: red (Hap01, $n = 1$ ), cyan (Hap02, $n = 1$ ), orange (Hap03, Hap04, $n = 2$ ), magenta (Hap05–Hap09, Hap18, $n = 6$ ), light violet (Hap10–Hap17, Hap19–Hap31, Hap33, Hap37, Hap52, Hap53, Hap58, Hap66, Hap70, $n = 28$ ), and light grey (Hap32, Hap34–Hap36, Hap38–Hap51, Hap54–Hap57, Hap59–Hap65, Hap67–Hap69, $n = 32$ ). AMOVA necessitates grouping haplotypes because ungrouped singletons yield zero variance estimates. ##Puerto Rico is categorized separately from the continental United States in this analysis, as the species was initially introduced to this US territory. This distinction reflects its role as the earliest point of establishment. For the continental analysis, all countries and territories were grouped into Asia and North America.

### 3.8 AMOVA analysis

Hierarchical AMOVA indicated that the majority of genetic variance was attributable to differences within haplogroups (σ² = 4.160 of 4.399 total), while only a small proportion was explained by variation among haplogroups (σ² = 0.238; ΦST = 0.05418, p < 0.001; Table 2). When sequences were grouped by host plant family, 6.61% of the total variance was accounted for (σ² = 0.291 of 4.399 total; ΦST = 0.06614, p = 0.004), suggesting a modest yet statistically significant association between mitochondrial diversity and host utilization. By contrast, country-level grouping explained only 1.99% of the variance (σ² = 0.087 of 4.399 total; ΦST = 0.01987, *p* = 0.061), and continental grouping contributed even less, with a negative variance component (σ² = –0.158 of 4.358 total; ΦST = 0.01067, *p* = 0.240). The non-significant country- and continent-level *Fst* values likely reflect uneven sampling, as only a handful of specimens were available from China, Hong Kong, Iraq, and the US. Consequently, genetic structure at these geographic scales could not be robustly assessed (Table 2).

### 3.9 Tajima’s test analysis

Analysis of 373 mitochondrial sequences (487 bp) identified 95 segregating sites, yielding a mutation-based diversity estimate of θ = 0.030023. Nucleotide diversity was low (π = 0.004009), corresponding to an average of 1.95 pairwise nucleotide differences across sequences. Tajima’s neutrality test produced a strongly negative value (D = –2.5566), indicating a significant excess of low-frequency polymorphisms. A complementary analysis of the 70 unique haplotypes recovered the same 95 segregating sites and 105 total mutations, with higher haplotype-level nucleotide diversity (π = 0.01140) and an average of 5.55 nucleotide differences (k). Mutation-based θ was similarly elevated (θ per site = 0.04474), and Tajima’s D remained strongly negative (D = –2.550, p < 0.001). The concordance between sequence-level and haplotype-level results consistently supports a demographic history characterized by recent population expansion following a bottleneck or the action of purifying selection, reducing intermediate-frequency variants.

### 3.10 Haplotype code correspondence across published studies

Direct comparison of published *mtCOI* datasets showed extensive overlap among haplotype codes assigned by different authors. Five *mtCOI* codes reported by Akmal et al. (2017) (PKDG4, PKJG1, PKLH2, PKMH2, and PKOK3) were congruent with AB4 in Kranthi et al. (2017), corresponding to Hap02 in our framework. PKSL2 from Akmal et al. (2017) aligned with PBBT76 in Kranthi et al. (2017), defining Hap08. PKBN1 matched haplotype 4 in Sagarbarria et al. (2020), which we classify as Hap37.

A larger set of 44 *mtCOI* haplotypes reported by Akmal et al. (2017) (PKBN2–5, PKBP2–4, 6, 8–13, PKDG3, PKKW1, 3, 5–8, 10–11, 14, 16, 18–19, 36, PKLD1, 3–4, 6, PKMH5, PKMN1–3, 6, PKOK4, PKSL4, PKTS1, and PKVR1, 6–8) corresponded directly to the 31 haplotypes described by Kranthi et al. (2017) (BHT, C4, FB12, 14, FZ13, FK107, NG4, PBBT38, 46, 50, 52, 54, 67, 80, 86, RKO, RJHG12, SS12, VDL, SG1, 3, 6–7, 15, 17, 20, 26, 32–33, and SGd11–12). These sequences also matched haplotypes 1 and 11 in Sagarbarria et al. (2020) and the 81 JAS variants reported by Sharma et al. (2025) (JAS1–5, 7–9, 10–34, 35–48, 49, 51–54, 56–60, 62–67, 68–72, and 73–85). In our phylogenetic reconstruction, all of these sequences formed a single, well-supported clade that we designate as Hap01.

Across studies, many *mtCOI* designations therefore represented the same underlying genetic lineages. To resolve these inconsistencies, we assembled a unified database containing all published *mtCOI* sequences together with previously unpublished records retrieved from GenBank and BOLD and standardized them into a consistent haplotype nomenclature.

## 4 DISCUSSION

Phylogenetic inferences from a single gene can lack resolution at broader taxonomic depths; however, for the specific aim of confirming whether our sequences and identified reference sequences from GenBank and BOLD form a single, well-supported clade, the *mtCOI* gene is especially suitable. The standard barcode fragment contains conserved primer-binding sites and sufficient internal variability to resolve closely related lineages (Folmer et al., 1994; Souza et al., 2016). Its substitution rate, an order of magnitude higher than that of most nuclear markers, is driven by limited mitochondrial DNA repair capacity, persistent oxidative damage within the organelle during adenosine triphosphate production, and the absence of histones (Li et al., 2019; Chial & Craig, 2008). Maternal inheritance without recombination preserves haplotype integrity, while the germline bottleneck during oogenesis intensifies genetic drift, expediting lineage sorting and the fixation of novel variants (Rebolledo-Jaramillo et al., 2014). These combined properties conserved primer sites, elevated mutation rate, minimal recombination, and the germline transmission bottleneck make *mtCOI* an optimal marker for verifying clade membership and resolving recent divergences.

Although truncated *mtCOI* fragments (e.g., under 500 bp) can potentially resolve intraspecific haplotypes, their performance appears highly context-dependent. For instance, Zhao et al. (2013) removed all sequences shorter than ∼400 bp, corresponding to a group I intron, from their Paramecium dataset, and yet recovered 90 distinct haplotypes, indicating that moderately trimmed coding regions may retain essential polymorphisms. Conversely, Souza et al. (2016) subdivided the standard 658 bp Folmer barcode into ∼650, ∼450, and ∼400 bp subfragments for *Coreidae* and *Pentatomidae*, observing fragment-dependent shifts in topology and reduced node support at deeper phylogenetic levels. By focusing on intraspecific analysis using a standardized 487 bp *mtCOI* segment, which is longer than the shortest fragments tested, we expect to preserve sufficient variation for haplotype-network reconstruction and basic population-genetic analysis, while potentially reducing biases associated with very short amplicons and avoiding the need for additional sequencing. It should be noted that limitations of shortened fragments may still arise at broader taxonomic scales, although these effects may be less pronounced in an intraspecific framework. Nonetheless, we recognize that methodological heterogeneity may introduce bias.

We therefore advocate for future collaborative efforts focused on resequencing under standardized protocols, incorporating additional loci, and expanding international sampling. Such initiatives will enable the development of a more comprehensive multilocus database, improving phylogenetic resolution and informing effective management strategies for this invasive pest. Crucially, invasive populations in the southeastern US share the Hap01 lineage on cotton, eggplant, hibiscus, and okra. Common Malvaceae hosts in the southeastern US including okra (*A. esculentus)*, hollyhock (*Alcea rosea*), and rose mallow (*Hibiscus moscheutos*), as well as hibiscus (*H. rosa-sinensis*), likely serve as refugia when cotton is absent, sustaining local *A. biguttula* populations between seasons. Additionally, the hibiscus species *H. syriacus* (Rose of Sharon), which is more common in northern states, may become susceptible to this pest as its range expands northward. Hap01’s ability to exploit diverse hosts spanning Malvaceae (okra, cotton, hibiscus, hollyhock), Solanaceae (eggplant, potato), Pedaliaceae (sesame), Fabaceae (soybean), and multiple ornamental genera underscores its ecological plasticity and its potential to overwinter and bridge seasons on non-cotton Malvaceae hosts in the US. These findings emphasize the need to include both vegetable and ornamental Malvaceae in quarantine monitoring and integrated pest management to disrupt continuous generational carryover. Haplotype 48, collected from eggplant in India and presumed to be correctly identified morphologically, given the common occurrence of this species in the region, is genetically divergent (up to 3.0%) from all other haplotypes, suggesting a potential cryptic species (i.e., morphologically indistinguishable but genetically distinct). In contrast, Haplotype 58, recovered from potato, an atypical host for this hopper in India, is also genetically divergent (up to 3.67%), raising the possibility of misidentification or sample mislabeling and indicating that it might represent a different *Amrasca* species. The divergence of Hap48 and Hap58 suggests that sequencing an additional gene, applying integrative taxonomy, and conducting detailed morphometric analyses could help clarify whether they represent long-term endemic lineages or potential cryptic species.

We characterized patterns of haplotype diversity and relative divergence within the *mtCOI* network, following standard interpretations of network topology (e.g., centrality, connectivity, and geographic breadth; Ahmed et al., 2024), rather than inferring explicit haplotype ages. Across the 70 haplotypes, Hap01 (red group) occupied the most central, frequent, and broadly distributed position, indicating low relative divergence and a major hub within the network. Hap02 (cyan group), along with Hap03 and Hap04 (orange group), formed the next set of closely connected haplotypes, each showing short mutational distances to Hap01.

Within the magenta group, Hap09, Hap07, and Hap05 were interior and more closely connected to the core, whereas Hap18, Hap08, and Hap06 were peripheral and more divergent, reflecting reduced connectivity. For the pink group, interior haplotypes (Hap31, Hap29, Hap28, Hap26, Hap25, Hap23, Hap21, Hap19, Hap17, Hap14) showed lower divergence from the central clusters. In contrast, peripheral haplotypes (Hap70, Hap66, Hap30, Hap27, Hap24, Hap22, Hap20, Hap16, Hap15, Hap13, Hap12, Hap11, Hap10) were more divergent and geographically restricted, primarily to India and Pakistan, except for Hap66 (Hong Kong). Several pink-group haplotypes (Hap52, Hap53, Hap37, Hap33) exhibited wider geographic ranges, indicating dispersal despite moderate divergence.

Grey group haplotypes were mostly singletons positioned at the network periphery, except for Hap42 and Hap35, which were more closely associated with Pakistani cotton samples. Highly divergent peripheral haplotypes included Hap65 (Iraq), Hap64–Hap62 (Bangladesh), Hap56–Hap50 (Philippines), and Hap66 (Hong Kong), each separated from the core clusters by multiple mutational steps. Hap58 (magenta), the only sequence labeled from potato, and Hap48 (grey), collected from eggplant, both fell outside the main network structure, suggesting possible misidentification or cryptic lineages; however, additional loci will be required to evaluate these possibilities.

Overall, genetic diversity was highest in South Asian populations (India, Bangladesh, and Pakistan), which contained both central and divergent peripheral haplotypes, whereas recently colonized or island populations (Hong Kong, the Philippines, and Puerto Rico) exhibited reduced diversity and predominantly peripheral haplotypes. Iraq showed an intermediate pattern, while China and the mainland US contained only Hap01, consistent with very recent introductions. Tajima’s D, calculated from all 70 unique haplotypes, was strongly negative (D = –2.55, ***P < 0.001), indicating an excess of rare variants consistent with population expansion or purifying selection.

Genetic patterns suggest that South Asia is the ancestral source of diversity. At the same time, island populations, such as those in Hong Kong, the Philippines, and Puerto Rico, represent introductions with reduced nucleotide diversity. The geographic pattern of haplotype diversity is consistent with either historical movement through trade or natural long-distance dispersal, but the present dataset does not allow us to distinguish among these mechanisms. South Asia, a major exporter of ornamentals and vegetables, provides frequent pathways for insect dispersal (Liebhold et al., 2012; Hulme, 2009). Hong Kong and the Philippines, long-standing maritime trade hubs that link South Asia with East Asia and the Pacific, exhibit genetic signatures of founder events followed by rapid expansion (McKeown, 2010). Puerto Rico functions as a Caribbean bridgehead into the Americas. Its low nucleotide diversity but high haplotype diversity is consistent with a bottleneck followed by local proliferation, a pattern also observed in other invasive insects introduced through Caribbean ports (Ahmed et al., 2024). The presence of Hap52 in both Puerto Rico and the Philippines suggests shared introduction history or secondary movement between these regions. At the same time, the discovery of Hap70 in Puerto Rico as a singleton points to an additional introduction from an unsampled or unknown source. The presence of both shared and singleton haplotypes in Puerto Rico suggests that more than one source population may be involved; however, the current dataset does not allow us to determine whether this reflects multiple introductions or a single introduction containing multiple haplotypes. The absence of unique haplotypes in the southeastern US, together with the presence of Puerto Rican haplotypes in that region, is consistent with Puerto Rico serving as the most likely source of the US population. This interpretation is further supported by the geographic proximity of Puerto Rico to the mainland US and by numerous documented cases in which insects have colonized the southeastern US via Caribbean islands (Hoddle, 2004; Keller et al., 2007).

Puerto Rico contains haplotypes also found in Asia and the Philippines, but the available data do not allow us to determine whether it serves as a regional source population or simply reflects limited sampling. Human-mediated movement through trade remains a plausible pathway for long-distance dispersal, but our data do not allow us to attribute specific introduction events to phytosanitary lapses. Instead, the results highlight the need for continued surveillance and improved diagnostic capacity to detect introductions early, regardless of pathway. In addition to human-mediated movement, long-distance atmospheric transport is a well-documented mechanism for dispersal in leafhoppers and other small hemipterans (Carlson et al., 1992). Prevailing winds, tropical storms, and monsoon systems can carry individuals over long distances, and such weather-driven dispersal could contribute to the presence of shared haplotypes across distant regions. This mechanism should be considered alongside trade-based pathways when interpreting invasion histories.

Integrative *mtCOI*-based studies in other invasive hemipterans highlight the value of consolidating publicly available sequence data into consensus haplotype frameworks, enhancing molecular identification, regulatory oversight, and informed management strategies. Ahmed et al. (2024) compiled existing *Thrips parvispinus* barcodes into a unified network, revealing a single dominant haplotype that drives the global spread and informs quarantine measures. In *Bemisia tabaci*, De Barro & Ahmed (2011) merged hundreds of *mtCOI* records to reconstruct global invasion pathways of cryptic species, while Ahmed et al. (2011) used consensus sequences and network analysis to distinguish sibling whitefly species in Egypt and Syria, enhancing diagnostic accuracy. Although NUMTs and other pseudogenes were later identified as sources of false-positive variation in some cryptic *B. tabaci* species, generating consensus sequences and applying haplotype network analysis, as demonstrated in these studies, has proven effective in resolving intraspecific variation in many studies. More recently, Peng et al. (2025) assembled worldwide *B. tabaci mtCOI* haplotypes to show that just three haplotypes account for the majority of invasions, thereby delineating putative native versus invaded ranges. These precedents underscore how consensus-haplotype databases not only clarify invasion histories but also underpin molecular tools, regulatory protocols, and targeted control strategies (Ahmed, 2025b). Our study on *A. biguttula* applies these principles: by curating 373 trimmed *mtCOI* sequences into 70 consensus haplotypes, we deliver a robust reference resource that will support future phylogeographic investigations, regulatory measurements, and management strategies for this pest. Our comprehensive analysis of all *mtCOI* sequences available in GenBank and BOLD revealed that the invasion of *A. biguttula* into the mainland US involved a single haplotype, mirroring patterns observed in other rapidly expanding invasive insects. For example, the global spread of the MEAM1 cryptic species in *B. tabaci* was driven by a single haplotype (De Barro & Ahmed, 2011). However, given that GenBank and BOLD entries derive from studies with heterogeneous sampling schemes across multiple laboratories in Asia and variable time frames, any conclusions regarding the putative native range, diversity inferences, or timing of invasion events should be interpreted with caution.

The *mtCOI* haplotypes reported by Akmal et al. (2017) largely overlap with those described by Kranthi et al. (2017) and Sagarbarria et al. (2020). In our phylogenetic reconstruction, these sequences cluster into a single, well-supported clade, which we designate Hap01. Multiple *mtCOI* naming schemes used across previous studies were found to refer to the same underlying lineage. For example, several haplotypes labeled differently in Akmal et al. (2017), Kranthi et al. (2017), Sagarbarria et al. (2020), and Sharma et al. (2025) all collapsed into a single, well-supported clade in our phylogenetic reconstruction. This demonstrates that much of the published variation represents synonymous labels for the same lineage, which we designate as Hap01. These patterns highlight the need for a unified, consensus haplotype framework to ensure consistent interpretation across studies.

The resulting repository now encompasses consensus haplotypes Hap01 through Hap70, representing the most exhaustive nomenclature of this species’ mitochondrial diversity to date. Each entry is annotated with precise geographic coordinates and host-plant associations, enabling integrative analyses of phylogeographic and host-specific dynamics. From these data, we infer that Hap01 occurs throughout the species’ putative native range and invaded regions and exploits a broad host spectrum, including cotton, ornamental plants, and vegetable crops. Although some singletons and shared haplotypes shown indication of geographic or host-plant fidelity, these patterns remain tentative and underscore the need for broader sampling and incorporation of additional genetic markers to validate and redefine these associations. Our consensus nomenclature, together with ecological metadata, provides a practical framework for future studies. It should simplify adding new *mtCOI* sequences and support analyses of evolutionary and ecological patterns across time and space.

These molecular insights, grounded in an *mtCOI*-derived, genetic identity–based framework that is indispensable for revealing new species (Ball & Armstrong, 2006), biotypes (Perring, 2001), cryptic species (Hebert et al., 2004), and haplotypes (Toda & Murai, 2007) beyond morphological resolution, link genetic variation to documented biology and ecology, refine delineation of native versus invaded ranges, and enable targeted regulatory and control strategies. The global known dispersal of *A. biguttula* appears influenced by horticultural trade (Mound & Collins, 2000), but long-distance atmospheric transport is also a documented mechanism for leafhopper movement, with prevailing winds and storm systems capable of carrying individuals over large distances (Carlson et al., 1992). Current regulatory frameworks and scientific resources remain insufficient to address the accelerating threat of invasive species (Hulme, 2021), underscoring the urgent need for user-friendly, field-deployable molecular diagnostics and standardized international phytosanitary protocols for early detection and mitigation.

## ACKNOWLEDGEMENTS

The authors gratefully acknowledge the students and staff at the Pee Dee Research and Education Center, Clemson University, Florence, South Carolina, USA, for their invaluable support. We thank Tariq Alam and Katherine Wakeley for expert assistance with molecular work. We are also grateful to the growers who allowed us to collect specimens from their crops. In addition, we thank Carmen Ketron, Matt Lennon, Powlomee Mondal, Shirley Cruz, Zachary Snipes, and Peilin Tan for their dedicated help with field collections. We are indebted to Lamb’s Produce and Plants (Florence, South Carolina, USA) for generously donating hibiscus plants, which were essential for maintaining our *Amrasca biguttula* colonies. Finally, we thank the anonymous reviewers for their constructive feedback, which helped improve the draft. Ahmed MZ acknowledges support from two multi-state projects, SC-1700692 (S-1073) and NCERA-224. Mention of trade names or commercial products in this publication is solely for the purpose of providing specific information and does not imply recommendation or endorsement by the USDA.

## AUTHORS’ CONTRIBUTIONS

All authors conceived the study. NY and SG conducted sequencing, while MZA conducted diagnostics, analyzed the data, and wrote the first draft. All authors reviewed, read, and approved the manuscript.

## DATA AVAILABILITY STATEMENT

The raw data used for the analyses are openly available in ‘Mendeley data.’ 10.17632/8ffrsf6ksf.1 (Ahmed, 2025).

## FIGURE LEGENDS

**SUPPLEMENTARY FIGURE 1 Supplementar Figure 1.**

Box-and-whisker plots of nucleotide diversity (π) per site across sampled populations. Boxes represent the interquartile range (IQR), horizontal lines indicate medians, whiskers extend to values within 1.5×IQR, and dots denote outliers from bootstrap replicates. “Overall” is retained for context but should be interpreted cautiously, as pooled distributions may obscure population-level structure. The USA shows near-zero π due to the fixation of Hap01. China is labeled “undefined” because the available sample size was insufficient to generate a distribution. Adequate sample sizes are required for meaningful bootstrap distributions. Kruskal–Wallis χ² and p-value are shown above the plot.

**SUPPLEMENTARY FIGURE 2** Median-joining networks (Leigh & Bryant, 2015) generated from all 373 sequences of *Amrasca biguttula* to illustrate haplotype relationships under three trait schemes: (a) overall network; (b) network colored by country of origin; (c) network colored by host plant family. Each hatch mark denotes a single mutational step, and each circle represents a unique haplotype, with circle size proportional to the number of individuals sharing that haplotype. The networks depict connections among haplotypes and the number of substitutions separating them (Templeton et al., 1992).

**SUPPLEMENTARY TABLE 1.** Partial *mtCOI* sequences of *Amrasca biguttula* retrieved from GenBank and the Barcode of Life Data Systems (BOLD), together with new sequences generated in this study. Sequences are annotated with host-plant records, geographic origin, haplotype assignments, and consensus sequences derived from the genetic analyses conducted herein.

**SUPPLEMENTARY TABLE 2.** Summary of *Amrasca biguttula* haplotype frequencies across host-plant species and geographical locations.

**SUPPLEMENTARY TABLE 3.** Haplotype frequencies in *Amrasca biguttula* populations.

**SUPPLEMENTARY TABLE 4.** Nucleotide diversity of *Amrasca biguttula* haplotypes by country of origin.

**SUPPLEMENTARY TABLE 5.** Genetic distances among *Amrasca biguttula* haplotypes.

