## Supplementary figures and images for "Global phylogeography of *Amrasca biguttula* (Hemiptera: Cicadellidae) across eight countries reveals a single-haplotype incursion into the United States beyond its putative native range"

### Figure S1.jpg

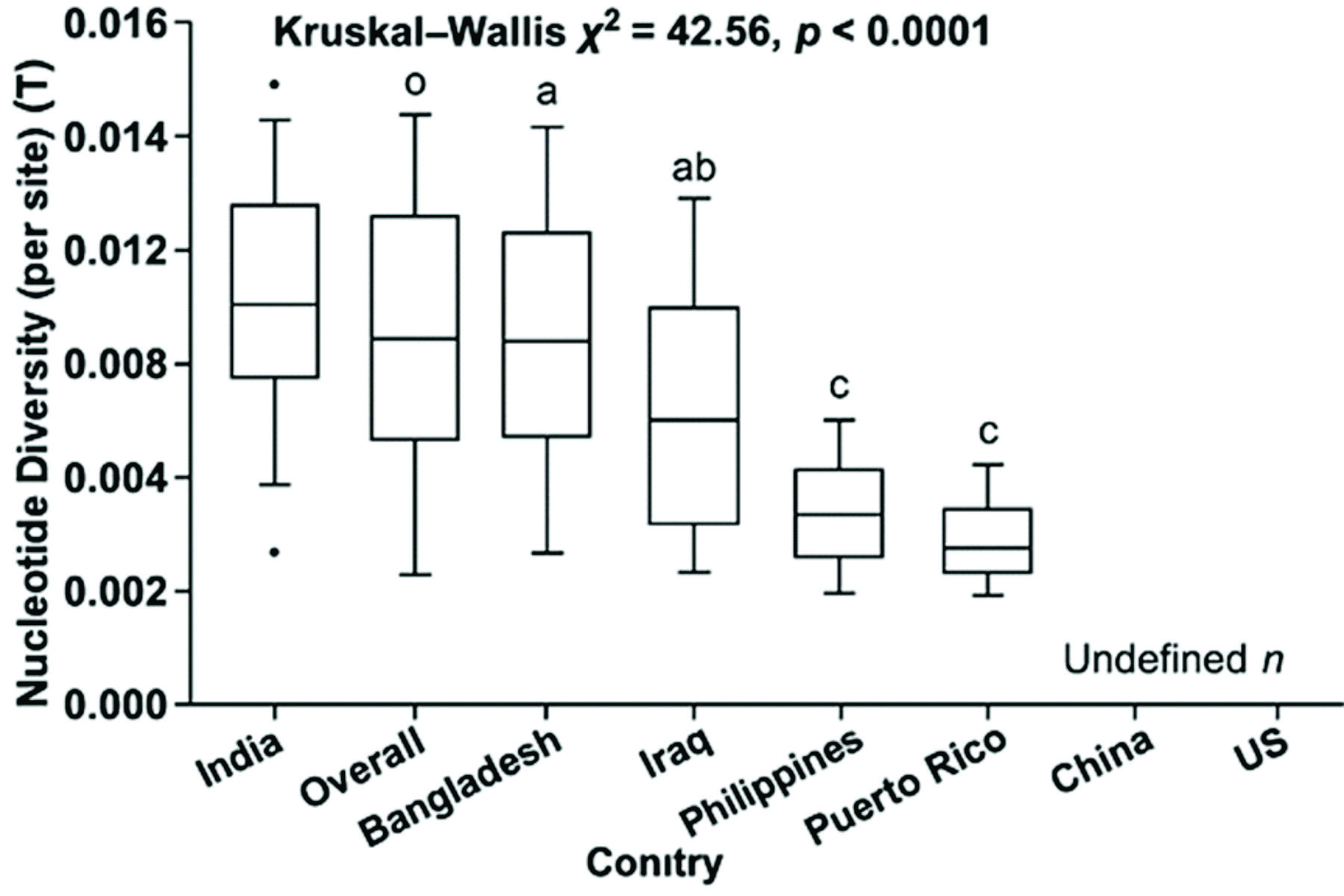

### Figure S2.jpg

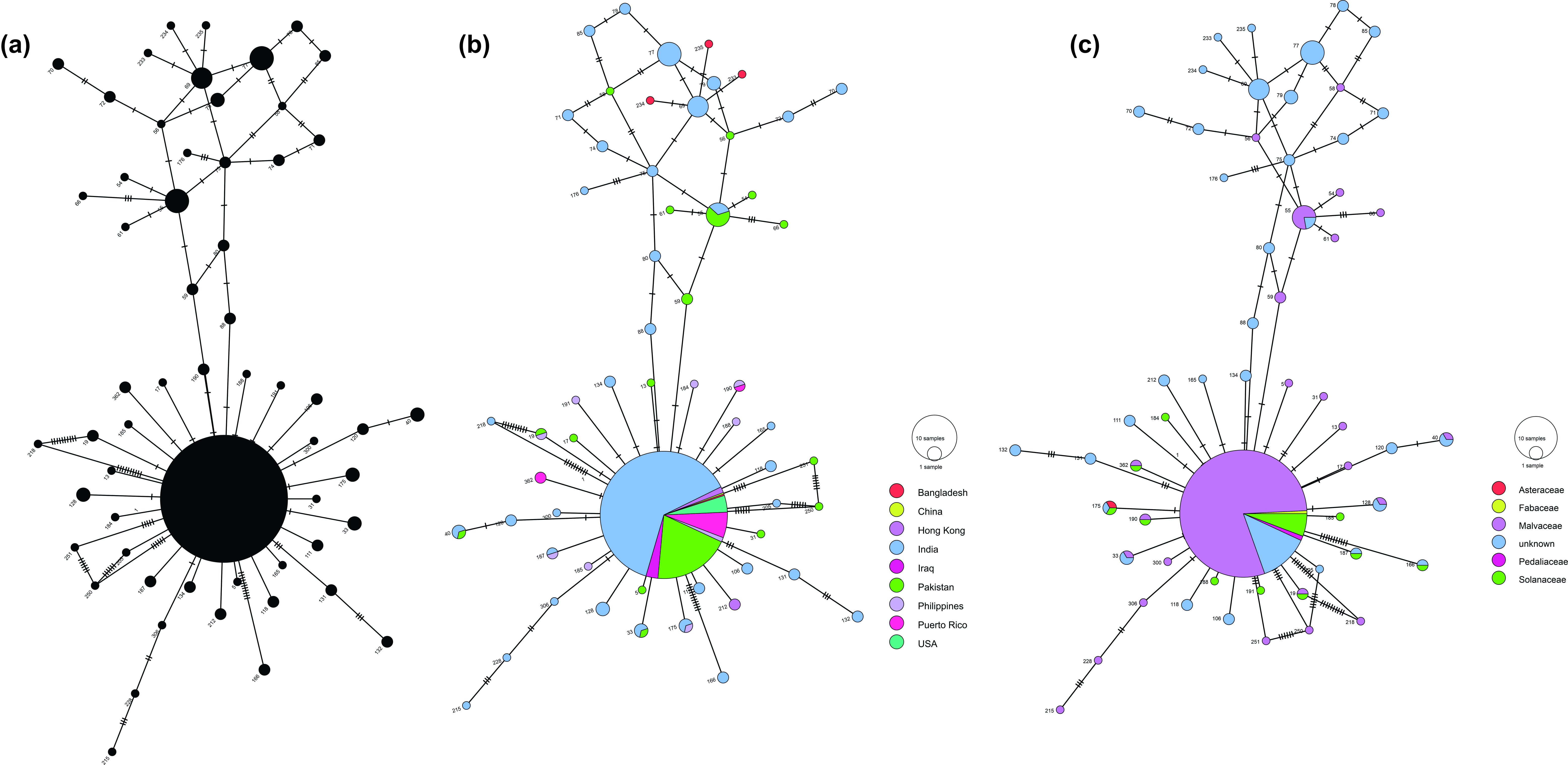
